# NGSEA: network-based gene set enrichment analysis for interpreting gene expression phenotypes with functional gene sets

**DOI:** 10.1101/636498

**Authors:** Heonjong Han, Sangyoung Lee, Insuk Lee

## Abstract

Gene set enrichment analysis (GSEA) is a popular tool to identify underlying biological processes in clinical samples using their gene expression phenotypes. GSEA measures the enrichment of annotated gene sets that represent biological processes for differentially expressed genes (DEGs) in clinical samples. GSEA may be suboptimal for functional gene sets, however, because DEGs from the expression dataset may not be functional genes *per se* but dysregulated genes perturbed by *bona fide* functional genes. To overcome this shortcoming, we developed network-based GSEA (NGSEA), which measures the enrichment score of functional gene sets using the expression difference of not only individual genes but also their neighbors in the functional network. We found that NGSEA outperformed GSEA in identifying pathway gene sets for matched gene expression phenotypes. We also observed that NGSEA substantially improved the ability to retrieve known anti-cancer drugs from patient-derived gene expression data using drug-target gene sets compared with another method, Connectivity Map. We also repurposed FDA-approved drugs using NGSEA and experimentally validated budesonide as a chemical with anti-cancer effects for colorectal cancer. We, therefore, expect that NGSEA will facilitate both pathway interpretation of gene expression phenotypes and anti-cancer drug repositioning. NGSEA is freely available at www.inetbio.org/ngsea.

## INTRODUCTION

Molecular phenotypes of clinical samples have proven useful in disease diagnosis, patient stratification, and drug discovery. Gene expression profiling is probably the most accessible strategy for molecular phenotyping of clinical samples. DNA chip technology and RNA sequencing have been widely used for molecular profiling of patient-derived primary cells and cell lines. Numerous gene expression profiles of clinical samples are now freely available from public data repositories such as the Gene Expression Omnibus (GEO) (Barrett *et al.*, 2013) and the National Cancer Institute Genomic Data Commons (NCI GDC) (Jensen *et al.*, 2017). Functional analysis of genome-wide expression phenotypes is generally more interpretable with annotated gene sets rather than individual genes; therefore, many bioinformatics methods for gene set analysis have been developed over the past several years (de Leeuw *et al.*, 2016). For clinical samples, the general purpose of gene set analysis of genome-wide expression profiles is to identify underlying disease-associated molecular processes, which can facilitate disease diagnosis and therapeutic intervention.

Two major approaches for gene set analysis of gene expression phenotypes are available: *over-representation* approaches and *aggregate score* approaches (Irizarry *et al.*, 2009). In the over-representation approach, a set of differentially expressed genes (DEGs) from the expression data set is selected, and then the significance of the over-representation of each annotated gene set among the selected DEGs is computed through a statistical test such as the hypergeometric test (Huang da *et al.*, 2009). This approach is reasonable but has some shortcomings (Pavlidis *et al.*, 2004; Irizarry, Wang et al., 2009). For example, in this approach, less-significant genes are treated as insignificant genes in the expression phenotype; the results, therefore, are highly dependent on the cutoff used for selecting DEGs. In addition, relative order information among the significant genes is not considered.

The analytical limitations of over-representation approaches can be overcome by aggregate score approaches, which assign scores to each annotated gene set based on all the gene-specific scores of the member genes. Gene set enrichment analysis (GSEA) (Subramanian *et al.*, 2005) is the most popular aggregate score approach available to date. In GSEA, genes for the expression profile are first rank-ordered by the gene-specific scores based on the expression difference, and then the enrichment score of each annotated gene set is computed based on a modified *Kolmogorov-Smirnov* (K-S) test. Despite its popularity, however, GSEA also has some shortcomings. For example, GSEA was designed to identify sets of genes that are differentially regulated in one direction, i.e., either up-regulated or down-regulated. If a gene set has matched genes for DEGs in which up-regulation and down-regulation are equally distributed, then its association with the expression phenotype may not be detected by GSEA. To overcome this limitation, a modified GSEA called absolute enrichment (AE) was developed that computes the absolute values of gene scores for both up- and down-regulated genes (Saxena *et al.*, 2006).

Another shortcoming of GSEA is that DEGs do not necessarily represent the functional genes that are responsible for the molecular processes represented by the gene sets. Instead, observed DEGs may be dysregulated genes perturbed by genuine functional genes in the molecular process of interest. Given that GSEA assigns a score to each gene set based on the scores of significant DEGs, a gene set comprising *bona fide* functional genes that exhibit no significant expression changes would not be captured by this method. This analytical limitation may be partially overcome by using annotated gene sets that are based on expression signatures rather than functional genes. For example, MSigDB, which was designed explicitly for use with GSEA, contains many signature gene sets derived from gene expression data (Liberzon *et al.*, 2011). The majority of databases of annotated genes for biological processes and diseases, however, are based on functional genes, such as disease-causing genes.

Network-based analysis of differential gene expression has been used to prioritize disease-causing genes (Nitsch *et al.*, 2009) and essential genes of cancer cell lines (Jiang *et al.*, 2015). These methods are based on the idea that functional genes for disease processes, such as tumorigenesis, tend to be surrounded by DEGs for that disease condition in the functional network. We, therefore, hypothesized that ordering genes by the differential expression of their local subnetworks (i.e., networks connecting each gene and its neighbors) will improve the ability to capture functional gene sets associated with the relevant biological processes. In this study, we present a network-based GSEA (NGSEA) that measures the enrichment scores of functional gene sets by utilizing the expression difference of not only individual genes but also their neighbors in the functional network. Although several network-based gene set analysis methods already have been proposed, these methods are modified from the over-representation approach, which identifies associations between two pre-selected gene sets, annotated gene sets from databases, or query gene sets from experiments based on relative closeness within the molecular network (Alexeyenko *et al.*, 2012; Glaab *et al.*, 2012; Wang *et al.*, 2012; McCormack *et al.*, 2013). To the best of our knowledge, NGSEA is the first network-based gene set analysis method that applies the aggregate score approach.

We found that NGSEA outperformed GSEA in retrieving KEGG pathway gene sets (Kanehisa *et al.*, 2017) for matched gene expression data sets. We also applied NGSEA to drug prioritization for several diseases and found that NGSEA performed substantially better than Connectivity Map (CMap) (Lamb *et al.*, 2006) in the ability to retrieve known drugs for matched cancer-associated gene expression data sets. We analyzed FDA-approved drugs to determine whether they had anti-cancer effects on colorectal cancer using NGSEA and experimentally validated the anti-cancer effect of budesonide, a chemical that is currently used as an anti-inflammatory drug. NGSEA is freely available for use as web-based software (www.inetbio.org/ngsea).

## MATERIALS AND METHODS

### Gene expression profiles, annotated gene sets, and a functional human gene network

To evaluate the gene set analysis performance for gene expression phenotypes, we used a gold-standard expression dataset composed of expression profiles in which their matched KEGG pathway terms are already annotated. We used KEGG disease datasets from GEO (KEGGdzPathwaysGEO) obtained from Bioconductor (https://bioconductor.org/packages/release/data/experiment/html/KEGGdzPathwaysGEO.html) as our gold-standard dataset to evaluate gene set enrichment analysis methods. This collection includes 24 expression data sets based on an AffyMetrix HG-U133a chip for which the phenotype is a disease with a corresponding pathway in the KEGG database (**Supplemental Table 1**). For example, the GSE21354 dataset of KEGGdzPathwaysGEO was annotated by the KEGG pathway term ‘glioma’ (hsa05214) and contains microarray-based gene expression data comprised of 14 samples from tumor tissues and four samples from normal tissues. These datasets have been previously used as a gold-standard dataset in comparing the performance of 16 gene set analysis methods (Tarca *et al.*, 2013).

We obtained pathway gene sets from human KEGG pathways (https://www.genome.jp/kegg/pathway.html as of June 2016) (Kanehisa, Furumichi et al., 2017) and drug-target gene sets from DSigDB version 1 (http://tanlab.ucdenver.edu/DSigDB/DSigDBv1.0/) (Yoo *et al.*, 2015) D1 data. Gene sets containing less than 15 genes were excluded from the analysis; this same criterion is used in the default parameter setting for GSEA. For data derived from DSigDB, drug names were mapped to the compound ID (CID) from the PubChem database (ftp://ftp.ncbi.nlm.nih.gov/pubchem/Compound/Extras/). A total of 276 KEGG pathway gene sets and 165 DSigDB gene sets were used in our final analysis. We used the following additional gene sets for web server construction: Gene Ontology biological process (GOBP) annotations (http://www.geneontology.org as of April 4, 2018) (Ashburner *et al.*, 2000), curated annotations of DisGeNET (http://www.DisGeNET.org as of June 8, 2018) (Pinero *et al.*, 2017), and disease gene annotations with more than three-star scores in DISEASES (https://diseases.jensenlab.org) (Pletscher-Frankild *et al.*, 2015).

To benchmark the ability to retrieve drugs for diseases, we compiled 17,063 links between 2,109 diseases and 1,481 chemicals based on a direct evidence of association as determined from the ‘therapeutic’ category of the Comparative Toxicogenomics Database (CTD) (http://ctdbase.org/ as of October 4, 2016) (Davis *et al.*, 2017). We combined information for drugs with synonyms in the CID.

For network-based analysis of the differential expression of genes, we employed a genome-scale functional gene network, HumanNet-EN (manuscript submitted), which is available from www.inetbio.org/humannet. Briefly, HumanNet-EN was constructed by integrating the functional associations between genes inferred not only from protein-protein interactions but also from diverse types of omics data using Bayesian statistics. The HumanNet-EN contains 424,501 functional links between 17,790 human genes (i.e., 94.6% of the coding genome). We also included a functional gene network for mouse, MouseNet (www.inetbio.org/mousenet), which contains 788,080 links between 17,714 mouse genes (i.e., 88% of the coding genome) (Kim *et al.*, 2016), to allow users to analyze mouse gene expression phenotypes with NGSEA.

### Running GSEA, AE, and NGSEA

We obtained the freely available software program javaGSEA version 3.0 from the Broad Institute (http://software.broadinstitute.org/gsea/downloads.jsp) and used it for the analyses and web server implementation. javaGSEA can analyze the input data as either a gene expression matrix (GSEA) or a pre-ranked list of genes (GSEA-preranked). The gene expression matrix needs to contain both control samples and case samples. One goal of our analysis was to improve GSEA by modifying the rank-order of genes; therefore, we used the GSEA-preranked function for the analyses in this study. In particular, we used ‘weighted GSEA-preranked’ with the default parameters. The original GSEA ranked genes based on either gene-based scores, the signal-to-noise ratio (*SNR*), or the log base 2 of the expression ratio (i.e., log_2_(Ratio)) from the most upregulated gene. *SNR* is calculated as the average expression value difference between the case samples and control samples divided by the sum of the standard deviations of each group of samples. The log_2_(Ratio) is computed by taking the logarithm (base 2) of the ratio between the average expression value of the case samples and the average expression value of the control samples.

In NGSEA, the original gene-based score was modified via network-based integration of the gene-based scores for network neighbors. We assigned a network-based score (*NS*) for each gene by integrating the absolute value of its gene-based score with the mean of the absolute value of the gene-based scores of its network neighbors using the following equation:

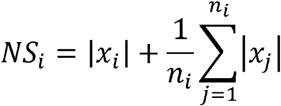

where *n*_*i*_ is the number of network neighbors of the *i*^*th*^ gene and *x*_*i*_ represents the score of the *i*^*th*^ gene. If a gene has no expression data, then we assigned the gene-based score to be zero. We initially tested both *SNR* and log_2_(Ratio), and found that the log_2_(Ratio) performed better in general; therefore, all the results presented in this report were based on the log_2_(Ratio). For the AE analysis, we ordered genes based on the absolute values of the log_2_(Ratio).

We performed GSEA, AE, and NGSEA with a gene list ordered by the log_2_(Ratio) value, the absolute value of the log_2_(Ratio), and the *NS*, respectively, using the GSEA-preranked function, from which we computed enrichment scores (*ES*), normalized enrichment scores (*NES*), *P*-values, and false discovery rate (*FDR*) values for each gene set based on a modified K-S test. To evaluate gene set recovery performance, we prioritized gene sets based on their absolute *NES*; this measure was used because gene sets with high scores for both positive and negative directions are equally weighted in GSEA.

### Drug repositioning using Connectivity Map (CMap)

We prioritized FDA-approved drugs for the 24 KEGG disease gene expression datasets using the CMap web server (https://portals.broadinstitute.org/cmap). CMap requires up and down tag lists (AffyMetrix HG-U133a probe ID) as input data; therefore, we selected the 50 highest up- and down-regulated probe IDs from each of the 24 disease expression data sets. If input genes were not based on AffyMetrix HG-U133a probe IDs, then we converted them to AffyMetrix HG-U133a probe IDs to run the CMap analysis.

### Anti-cancer activity analysis using a cell viability assay

We conducted MTS (3-(4,5-dimethylthiazol-2-yl)-5-(3-carboxymethoxyphenyl)-2-(4-sulfophenyl)-2H-tetrazolium) assays to measure cell viability following drug treatment. We used two colorectal cancer cell lines, HCT116 and HT-29, which were obtained from the Korean Cell Line Bank, for the assay. We purchased two candidate drugs, dobutamine, and budesonide, from Sigma (St. Louis, MO, USA). We dissolved chemicals in dimethyl sulfoxide (DMSO) prior to treatment. Cells were treated with the candidate drugs at concentrations ranging from 50 to 250 μM for 24, 48, and 72 hours. MTS reagents then were added to the cells. The number of viable cells was counted based on the absorbance at 490 nm on an ELISA microplate reader (Molecular Devices, San Jose, CA, USA), and the cell viability percentage was calculated. All experiments were repeated six times.

## RESULTS

### Overview of NGSEA

As summarized in **Figure 1**, NGSEA differs from GSEA in the method of scoring genes, resulting in different gene orders between the two methods. The list of genes generated by GSEA is ordered by the log_2_(Ratio) gene score. In contrast, the list of genes generated by NGSEA is ordered using a network-based score. This score was based on two assumptions. The first assumption is that the annotated gene set for biological processes that are truly associated may contain both up- and down-regulated genes. Whereas GSEA was designed to find gene sets that are regulated in one direction, groups of genes or systems are often regulated in both directions. To address this problem, NGSEA uses the absolute value of the log_2_(Ratio) for analyses; a similar approach was employed previously in an AE analysis (Saxena, Orgill et al., 2006). The second assumption is that the expression perturbation of gene regulators may cause severe dysregulation of their downstream genes such that the functional importance of a regulator for a given biological context would, in fact, be much greater than estimated by its own expression change. Thus, we expected that the expression difference in the local subnetwork would assign higher scores than the original gene-based score to truly functional genes. To address this problem, NGSEA integrated the mean of the absolute value of the log_2_(Ratio) for the network neighbors of each gene to account for the regulatory influence on its local subsystem.

**Figure 1.**
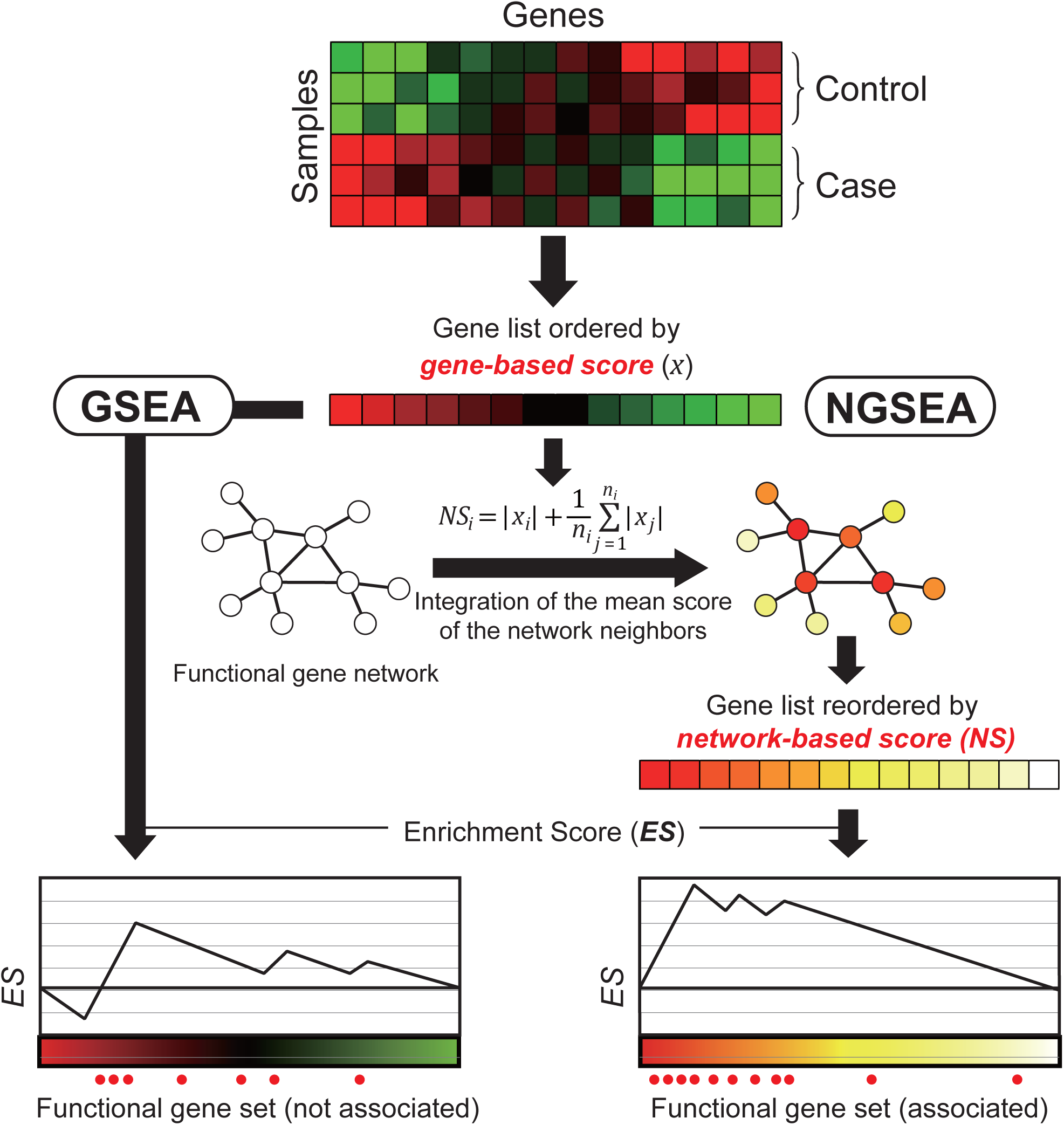
Overview of NGSEA. In the original gene set enrichment analysis (GSEA), genes of a given expression dataset are ordered by gene-based scores (e.g., signal-to-noise ratio [*SNR*] or log_2_(Ratio)) based on the gene expression difference between control and case samples. In network-based GSEA (NGSEA), the genes are ordered by network-based scores, which integrates the gene-based score with the mean of the scores of its neighbors in the functional gene network. This method is based on the observation that functional genes tend to cause expression changes in their network neighbors and therefore are more likely to be correlated with network-based scores.

### NGSEA outperformed GSEA and AE in identifying KEGG pathways for matched disease expression data sets

We evaluated the ability of GSEA, AE, and NGSEA to retrieve annotated gene sets for matched gene expression data sets. For this analysis, we used gold-standard gene expression datasets from the Bioconductor’s KEGGdzPathwaysGEO package. We used a total of 24 expression data sets associated with 12 different diseases (**Supplemental Table 1**), which were previously used as gold-standard datasets in comparing the performance of 16 gene set analysis methods (Tarca, Bhatti et al., 2013). We scored 276 human KEGG pathway gene sets (Kanehisa, Furumichi et al., 2017) that contained more than 15 member genes for each of the 24 expression datasets with GSEA, AE, and NGSEA, with an aim of retrieving the associated KEGG pathway term for each disease expression data set within the top predictions.

We observed a significantly higher rank distribution using NGSEA compared with GSEA and AE (*P*=2.35e-3 and *P*=4.0e-3, respectively, by Wilcoxon signed-rank test) (**Figure 2A**). The ranks for matched KEGG pathway terms were improved in NGSEA compared with GSEA in 18 of the 24 (75%) tested disease expression datasets (**Figure 2B**). For example, the KEGG pathway term for ‘Glioma’ was ranked as 131 via GSEA but as 18 by NGSEA for a gene expression dataset derived from glioma samples (GSE21354). Notably, the performance of the AE method was not significantly improved from GSEA (*P*=0.11 by Wilcoxon signed-rank test). These results clearly indicate that the major factor contributing to the improvement in NGSEA was the network-based analysis of the gene expression data.

**Figure 2.**
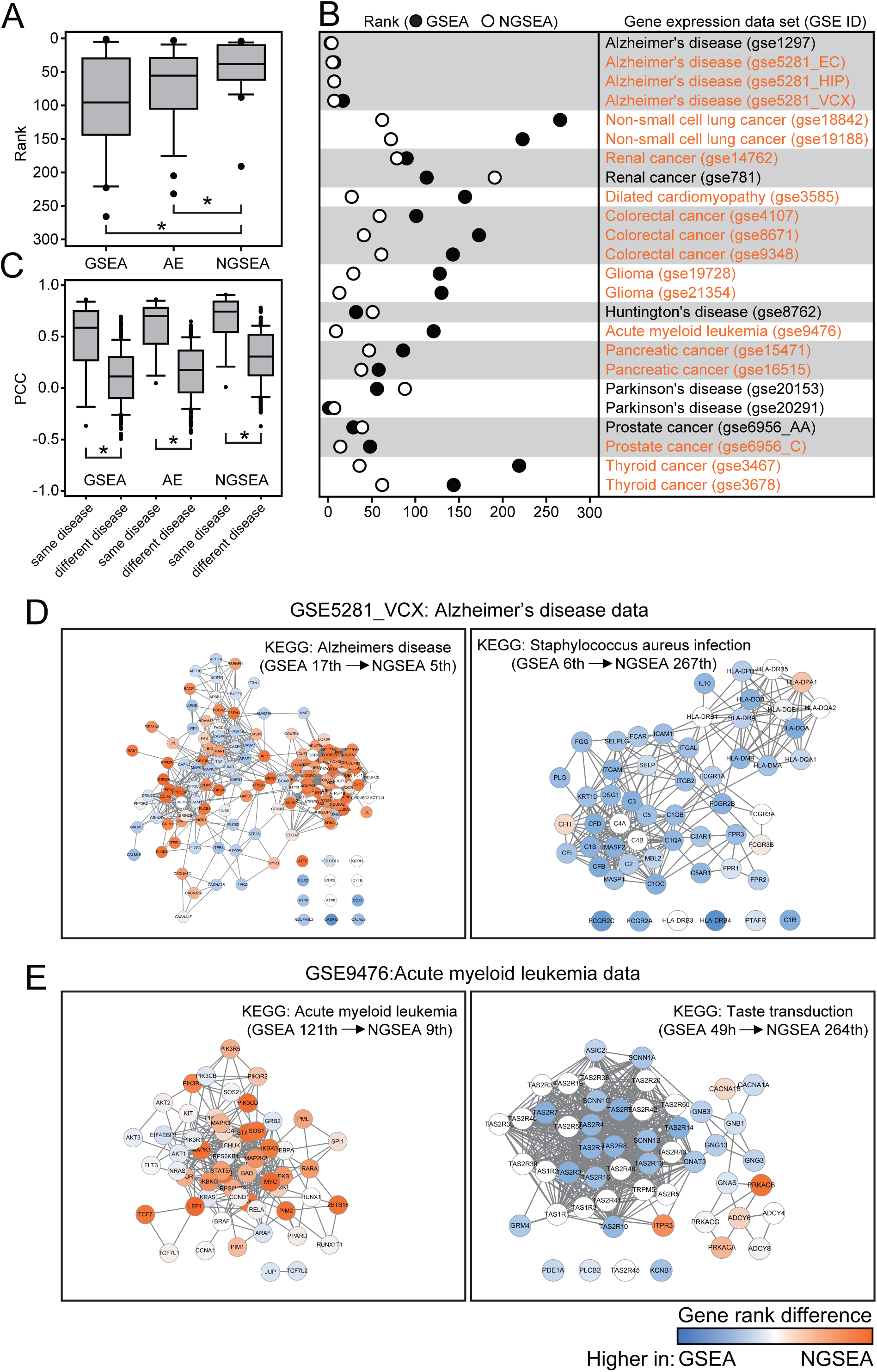
Recovery of KEGG pathways for matched disease expression datasets by GSEA, AE, and NGSEA. (A) Rank distribution of the matched KEGG pathway terms (out of 273 terms in total) for each of 24 gold-standard expression datasets from KEGGdzPathwaysGEO using gene set enrichment analysis (GSEA), absolute enrichment (AE), and network-based GSEA (NGSEA). The significance of the difference in the rank distribution was assessed by Wilcoxon signed-rank test (asterisk (*) indicates *P* < 0.05). (B) Rank comparison of the matched KEGG pathway terms between GSEA and NGSEA for each of the 24 gold-standard expression data sets. (C) Distribution of the Pearson’s correlation coefficient (*PCC*) of the normalized enrichment scores (*NES*) between the same diseases and different diseases. The significance of the difference in the rank distributions was assessed by Wilcoxon rank-sum test (asterisk (*) indicates *P* < 0.05). (D) Subnetworks for the KEGG pathway terms ‘Alzheimer’s disease’ (HSA05010) and ‘Staphylococcus aureus infection’ (HSA05150). The difference between the ranks assigned by GSEA and NGSEA is indicated by the color code (red and blue for higher ranking by NGSEA and GSEA, respectively) for each pathway member gene. (E) Subnetworks for the KEGG pathway terms ‘acute myeloid leukemia’ (HSA05221) and ‘taste transduction’ (HSA04742). The color code is the same as in (D).

Next, we tested the robustness of the three enrichment analysis methods by comparing the assigned scores for the KEGG pathway terms between the different expression profiles for the same disease. The 24 expression data sets were derived from 12 diseases, and nine of the diseases have multiple expression data sets. We hypothesized that if an enrichment analysis retrieved pathways based on disease-specific signals rather than technical variation, then scores for the pathways between two different expression datasets for the same disease should have a higher correlation than those for different diseases. We, therefore, computed Pearson’s correlation coefficients (*PCC*) between expression data sets using the normalized enrichment scores (*NES*) for all the test KEGG pathways. Then, we compared the distributions of *PCC* values between the same diseases or between different diseases. As expected, higher correlations were observed between the same diseases compared with different diseases for all three enrichment analyses (**Figure 2C**). Notably, we observed an improvement in the significance of the correlations difference between the same disease groups and different disease groups using NGSEA compared with GSEA (*P*=2.72e-6 and *P*=3.44e-5, respectively, by Wilcoxon rank-sum test). These results suggest that the enrichment analysis conducted using NGSEA may be less affected by variation among expression profiles for the same disease processes.

As expected, the improved ranks for the matched KEGG pathways were due to improved ranks of their member genes in the gene list used for the enrichment analysis. For example, the network-based scoring method improved the rank of the KEGG term ‘Alzheimer’s disease’ from 17^th^ to 5^th^ and reduced the rank of an irrelevant KEGG term ‘Staphylococcus aureus infection’ from 6^th^ to 267^th^ for the gene expression data set for Alzheimer’s disease (GSE5281_VCX). The majority of relevant pathway genes were ranked higher by NGSEA (red color) compared with GSEA (**Figure 2D**). As another example, we observed similar trends in the rank changes between relevant and irrelevant pathway terms for the KEGG term ‘acute myeloid leukemia’ (**Figure 2E**). These results demonstrate that the use of network-based scoring in enrichment analysis increases the ranks of truly functional genes within the ordered gene list, resulting in the assignment of higher scores to gene sets truly associated with the underlying biological process.

### Application of NGSEA to drug-target gene sets improved retrieval of known drugs compared with Connectivity Map

GSEA is the algorithmic foundation of the most popular drug repositioning system, the Connectivity Map (CMap) (Lamb, Crawford et al., 2006). The previous CMap database, which was based on the AffyMetrix HG-U133a chip, contained more than 7,000 expression profiles representing 1,309 compounds. A recent release of CMap included a database of reference expression profiles with more than 1000-fold scale-up based on L1000 platform technology, which is a low-cost, high-throughput reduced representation expression profile method (Subramanian *et al.*, 2017). The CMap system prioritizes drugs for diseases based on an inverse relationship between disease expression profiles and drug treatment expression profiles. To conduct a web-based CMap analysis, users submit signature genes for a given disease (e.g., the 50 most up-regulated and 50 most down-regulated genes from the disease-associated gene expression dataset). GSEA is applied to assign scores to each drug based on the anti-correlation of disease signature genes with the genes ordered by expression changes in drug condition, which represents the strength of the drug response. In contrast to CMap, which uses expression data from drug-treated samples, the NGSEA-based drug prioritization method uses functional genes for the drug’s mode of action; target genes. We used target genes for each FDA-approved drug as functional gene sets to test the association to diseases based on a list of genes ordered by network-based scores computed from disease-associated expression data (**Figure 3A**). We compiled target gene sets for drugs from drug-target links based on active bioassays from the Drug Signature Database (DSigDB) (Yoo, Shin et al., 2015). We performed drug prioritization using NGSEA on 24 gene expression datasets for 12 diseases from KEGGdzPathwaysGEO and target gene sets for 165 FDA-approved drugs (i.e., those with more than 15 targets) from DSigDB.

**Figure 3.**
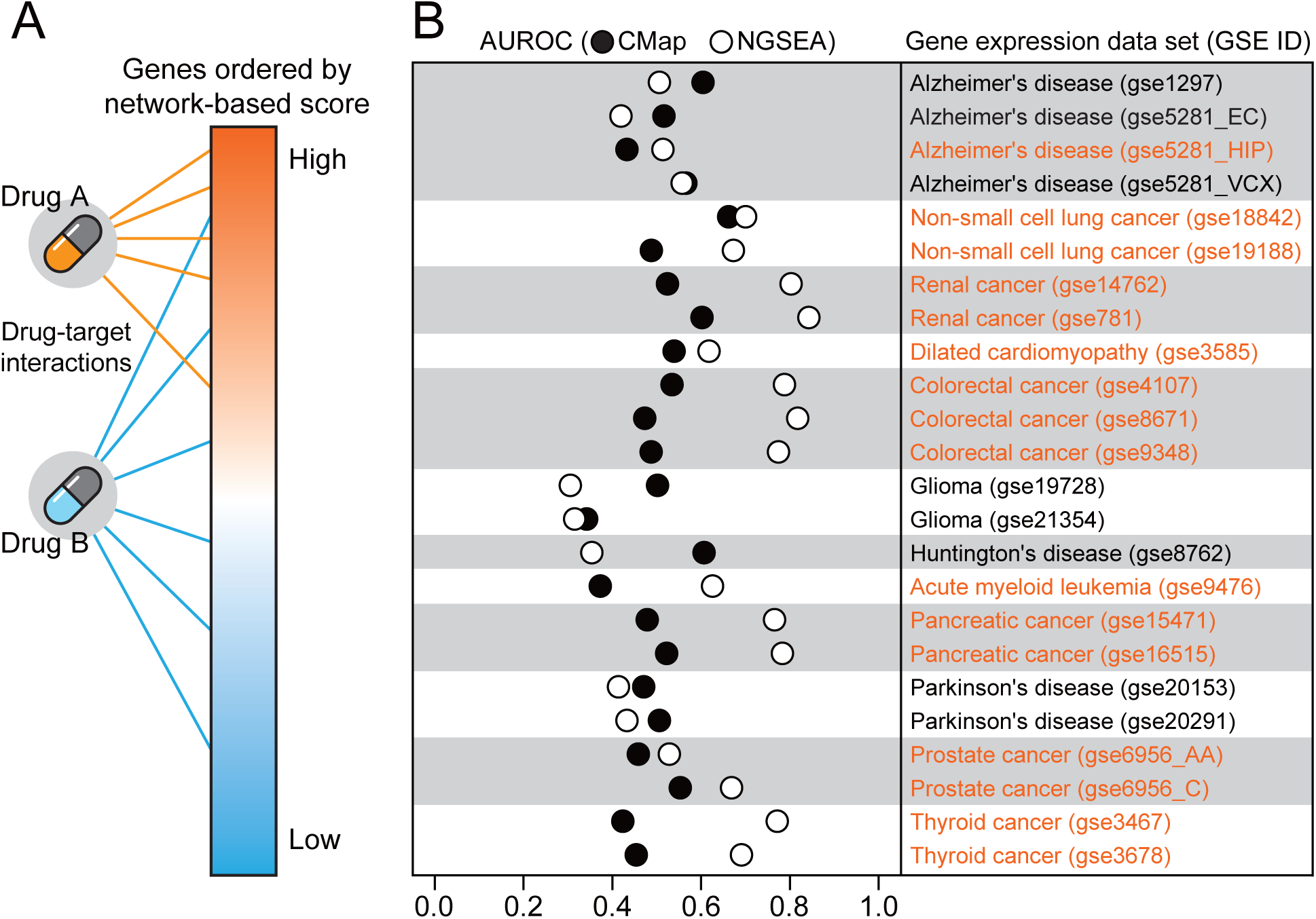
Retrieval of known drugs for matched disease expression datasets using CMap and NGSEA. (A) Overview of how network-based gene set enrichment analysis (NGSEA) retrieves drugs for the matched disease of the given gene expression data. Drug A targets genes with high network-based scores, whereas Drug B targets genes with a wide range of network-based scores; therefore, Drug A but not Drug B will be highly ranked. If Drug A but not Drug B is a known drug for the given disease, the prediction performance of NGSEA, as measured by the area under the receiver operating characteristic curve (AUROC) will be high. (B) Comparison of the AUROC between Connectivity Map (CMap) and NGSEA for the ability to retrieve known drugs for the matched disease of each gene expression data set.

We compared the ability of CMap and NGSEA to retrieve known drugs for each of the 24 disease-associated gene expression data sets. For benchmarking, we compiled 17,063 associations between 2,109 diseases and 1,481 chemicals based on direct evidence of an association from the ‘therapeutic’ category of the CTD (Davis, Grondin et al., 2017). The performance of both CMap and NGSEA were determined using the area under the receiver operating characteristic curve (AUROC). To avoid a biased evaluation due to differences in the number of drugs tested, we included only drugs that were considered in both CMap and NGSEA for the AUROC analysis. We found significantly improved AUROC for drug recovery using NGSEA compared with CMap (*P*=9.62e-4 by Wilcoxon signed-rank test). Recovery of known drugs was improved in 16 of 24 (67%) tested expression datasets using NGSEA compared with CMap (**Figure 3B**). NGSEA was particularly effective for retrieving cancer drugs; we observed improved performance in 14 of 16 (87.5%) cancer-associated expression datasets using NGSEA compared with CMap. These results suggest that using NGSEA with drug-target information may be an effective approach for anti-cancer drug repositioning.

### Identification of budesonide as a novel drug candidate to treat colorectal cancer

The effective retrieval of known drugs for various types of cancers suggested that we would be able to identify novel anti-cancer drugs using NGSEA. NGSEA yielded the highest improvement in the recovery of known anti-cancer drugs for colorectal cancer (GSE9348: AUROC=0.488 and 0.775 by CMap and NGSEA, respectively) (**Figure 4A**). We, therefore considered it highly likely that NGSEA would be able to identify novel drugs to treat colorectal cancer among the top repurposed FDA-approved chemicals. Among the top 30 chemicals predicted as candidates for colorectal cancer by NGSEA, six chemicals were currently used for the treatment of colorectal cancer and three additional chemicals had been tested in clinical trials for colorectal cancer (https://clinicaltrials.gov/) (**Figure 4B**). We also found evidence in the literature (via manual examination of the PubMed database) for anti-cancer effects in colorectal cancer for 13 additional chemicals predicted by NGSEA. We, therefore, considered the remaining eight candidates, for which there was no prior evidence of an anti-cancer effect in colorectal cancer, for follow-up experimental validation. Among the commercially available and affordable chemicals on our candidate list, we were able to obtain and test both dobutamine (5^th^) and budesonide (17^th^). We observed that budesonide significantly inhibited cell growth in two different colorectal cancer cell lines (HCT116 and HT-29) (**Figure 4C-D** and **Supplemental Table 2**). Budesonide was ranked 37^th^ by CMap compared with 17^th^ by NGSEA, suggesting budesonide would not be identified as a treatment for colorectal cancer using existing drug-repositioning analyses.

**Figure 4.**
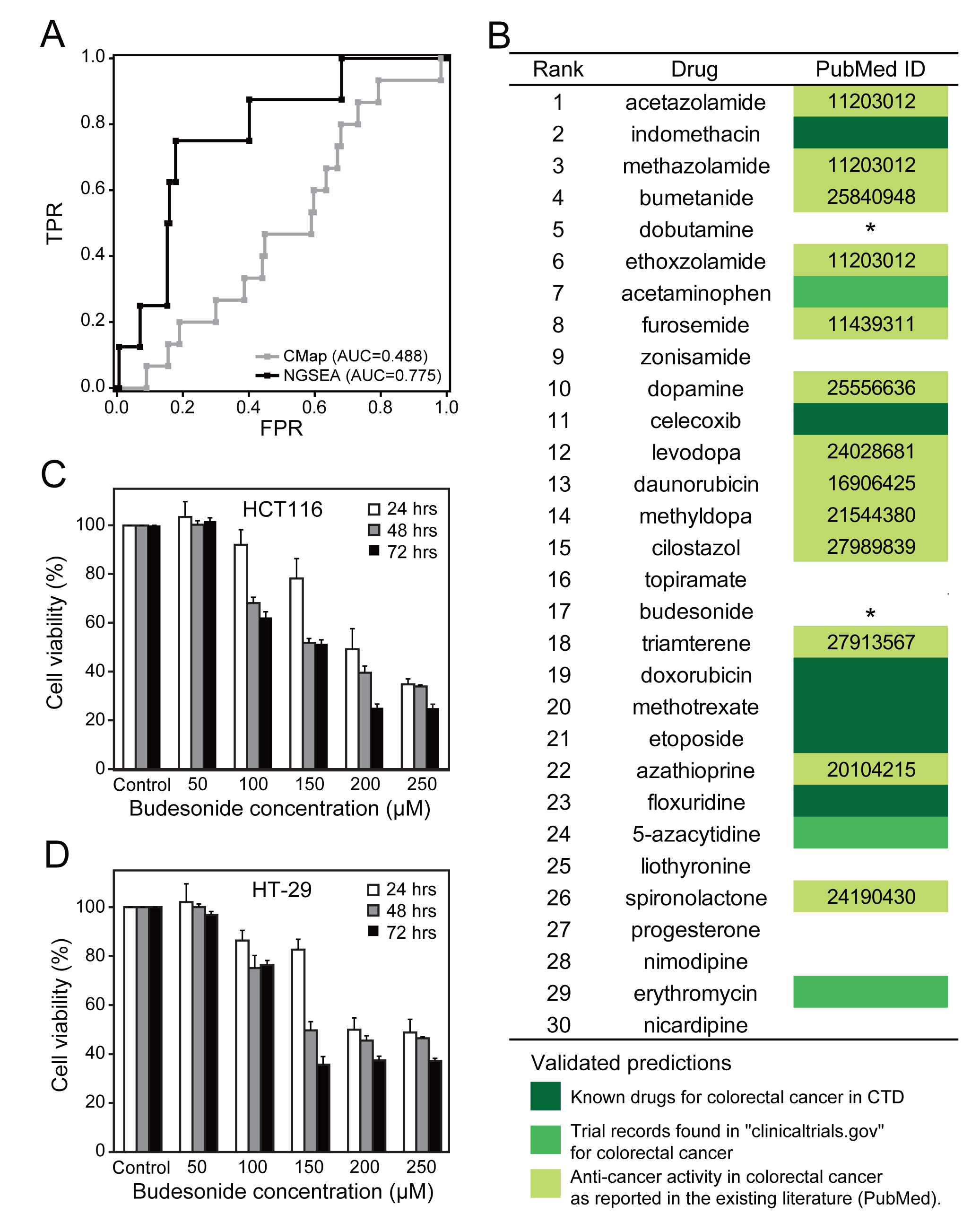
(A) Comparison of receiver operating characteristic (ROC) curves between Connectivity Map (CMap) and network-based gene set enrichment analysis (NGSEA) for the ability to retrieve known drugs to treat colorectal cancer using a gene expression data set from patients with colorectal cancer (GSE9348). (B) The top 30 chemicals for colorectal cancer as predicted by NGSEA. The color code indicates how the anti-cancer effect on colorectal cancer was validated for each predicted chemical. The asterisk (*) indicates the two drugs tested in this study. (C) Cell viability in the HCT-116 cell line after treatment with various concentrations of budesonide. (D) Cell viability in the HT-29 cell line after treatment with various concentrations of budesonide.

### Development of the NGSEA web server

To increase the usability of NGSEA, we have developed a web-based gene set enrichment analysis server (www.inetbio.org/ngsea). Users can prioritize functional gene sets representing biological and disease processes using various databases, including KEGG pathway (Kanehisa, Furumichi et al., 2017), GO biological process (Ashburner, Ball et al., 2000), DisGeNET (Pinero, Bravo et al., 2017), and DISEASES (Pletscher-Frankild, Palleja et al., 2015). Users can perform both GSEA and NGSEA simultaneously by submitting the gene expression phenotype. Both expression matrix (.gct format) data and the pre-scored list of genes (.rnk format) are allowed to be submitted as input data for the analysis. The default analysis runs for human genes, but enrichment analysis is also available for mouse genes using a genome-scale mouse functional gene network (Kim, Hwang et al., 2016; Kim and Lee, 2017). Users also can prioritize the gene sets by *ES, NES*, and *FDR*. Enrichment plots also are provided as output.

## DISCUSSION

In this report, we presented a network-based gene set enrichment analysis, NGSEA, which modified existing gene scoring methods by incorporating the differential expression information from neighbors in the functional gene network. We then demonstrated that NGSEA outperformed GSEA in retrieving both KEGG pathway terms and drugs for matched disease-associated gene expression data sets. Based on the benchmarking results, we have concluded that NGSEA will provide reliable functional information for interpretation of gene expression phenotypes of clinical samples. Most importantly, NGSEA performed well with functional gene sets. Because the original GSEA was designed to detect relationships between biological processes and chemicals based on expression signature genes, which are not necessarily functional genes, GSEA exhibited suboptimal performance with functional gene sets. MSigDB was designed explicitly to provide gene sets, many of which were derived from expression signatures, for GSEA. There are many other public databases, however, that contain annotated gene sets for diseases and pathways, and the majority of these gene sets include functional genes. We expect that NGSEA will be a useful tool for utilizing these available resources.

There are several existing methods that combine networks and gene set analysis. These methods measure the network distance between two gene sets (Alexeyenko, Lee et al., 2012; Glaab, Baudot et al., 2012; Wang, Hwang et al., 2012; McCormack, Frings et al., 2013). Although the sensitivity to detect the relationship between two gene sets was successfully improved using these methods compared with over-representation approaches, these methods still require the user to pre-select the query gene set by applying what is often an arbitrary differential expression score cutoff. These over-representation approaches to gene set analysis with network-based modification are therefore still limited by a lack of information regarding the relative orders between differentially expressed genes. We hypothesized that applying the network-based modification to an aggregate score approach to gene set analysis, which uses the differential expression values for the gene set analysis, would further improve the sensitivity to detect the relationship between two gene sets. To the best of our knowledge, NGSEA is the first method that combines network-based gene set analysis with an aggregate score approach.

We demonstrated that NGSEA could effectively retrieve known anti-cancer drugs for matched gene expression data sets. It is not clear why drug recovery using NGSEA was highly effective for different cancer types but not for neurodegenerative diseases such as Alzheimer’s disease, Parkinson’s disease, or Huntington’s disease (**Figure 3B**). One possibility is that the current human gene network model is short of predictive power for brain cells because we also observed low performance of NGSEA for brain cancer. We observed similar results using another high-quality genome-scale human gene network, STRING (Szklarczyk *et al.*, 2017). These findings suggest that we need to improve human functional gene networks before NGSEA can be used effectively for certain applications. Nevertheless, our results show that NGSEA is an effective tool for repurposing chemicals as anti-cancer drugs with the current functional gene networks.

## ACKNOWLEDGEMENTS

This work was supported by the National Research Foundation of Korea (NRF) grant funded by the Korean Government (MSIT) (NRF-2018M3C9A5064709, NRF-2018R1A5A2025079 to I.L.) to I.L.

## Conflict of Interest

none declared.

## Notes

https://www.inetbio.org/ngsea

